# Liquid Transmission Electron Microscopy for Probing Collagen Biomineralization

**DOI:** 10.1101/2023.06.19.545614

**Authors:** Liza-Anastasia DiCecco, Ruixin Gao, Jennifer L. Gray, Deborah F. Kelly, Eli D. Sone, Kathryn Grandfield

**Affiliations:** Department of Materials Science and Engineering, McMaster University, Hamilton, ON, Canada; Department of Biomedical Engineering, Pennsylvania State University, University Park, PA, USA; Institute of Biomedical Engineering, University of Toronto, Toronto, ON, Canada; Materials Research Institute, Pennsylvania State University, University Park, PA, USA; Center for Structural Oncology, Pennsylvania State University, University Park, PA, USA; Department of Materials Science and Engineering, University of Toronto, Toronto, ON, Canada; Faculty of Dentistry, University of Toronto, Toronto, ON, Canada; School of Biomedical Engineering, McMaster University, Hamilton, ON, Canada

**Keywords:** Liquid Electron Microscopy, Collagen Mineralization, Calcium Phosphate, Polyaspartic Acid, Hard Tissues

## Abstract

Collagen biomineralization is foundational to hard tissue assembly. While studied extensively, collagen mineralization processes are not fully understood as the majority of theories are derived from electron microscopy (EM) in static, dehydrated, or frozen conditions, unlike the liquid phase environment where mineralization occurs dynamically. Herein, novel liquid transmission EM (TEM) strategies are presented, where collagen mineralization was explored in liquid conditions for the first time. Custom thin-film enclosures were employed to visualize the mineralization of reconstituted collagen fibrils in a calcium-phosphate and polyaspartic acid solution to promote intrafibrillar mineralization. TEM highlighted that at early time points, precursor mineral particles attached to collagen and progressed to crystalline mineral platelets aligned with fibrils at later time points. This aligns with observations from other techniques and validates this liquid TEM approach. This work provides a new liquid imaging approach for exploring collagen biomineralization, advancing toward understanding disease pathogenesis and remineralization strategies for hard tissues.

## Liquid Transmission Electron Microscopy for Probing Collagen Mineralization

Biomineralization is key to the formation and maintenance of mineralized tissues found in the musculoskeletal system of vertebrates, such as bones and teeth.^1^ Bone is a biocomposite made primarily of calcium phosphate (CaP) mineral phase, known as hydroxyapatite, an organic phase consisting mainly of type I collagen and non-collagenous proteins (NCPs), and water.^2^ Notably, the distribution and extent of CaP among collagen fibrils are fundamental to the mechanical strength of mineralized tissues.^3,4^ Understanding the pathways behind collagen mineralization in tissues is thus significant for medical advancements; both to shed light on disease pathogenesis, such as periodontitis and osteoporosis, and to provide new insights for designing regenerative materials and treatment pathways.^5–7^

Although collagen biomineralization has been studied extensively,^8–14^ it is yet to be fully understood and well-controlled synthetically. For tissue regeneration applications, imitating the complex biomineralization behaviour of native collagen is key for engineering synthetic biomaterial models of similar structure, composition, and mechanical properties to facilitate tissue restoration.^7^ This has been achieved *in vitro* with the use of organic macromolecules to replace the role of NCPs in templating mineral formation.^9^ One such additive includes polyaspartic acid (pAsp), a synthetic mimic of NCPs believed to facilitate intrafibrillar collagen mineralization found in native mineralized tissues.^15,16^ pAsp is an established additive in *in vitro* collagen mineralization models, where its highly disordered structure is theorized to act as a nucleation inhibitor of CaP in solution.^9,17,18^ Several theories exist on the mechanisms behind how pAsp interacts with mineral ions and lead to collagen mineralization,^18–21^ where research into these nanoscale processes has been thus far limited by the technologies available to probe them directly.

To date, traditional and cryogenic (cryo) transmission electron microscopy (TEM) modalities have dominated the characterization of nanoscale collagen mineralization processes and helped elicit crucial theories of their mechanistic pathways.^9,11,20,22–28^ This research has led to tremendous strides within the collagen mineralization field, specifically involving the discovery of non-traditional CaP crystallization theories involving precursor phases unlike traditional ion-by-ion pathways.^28^ However, these TEM methods cannot visualize structures in the liquid state, representative of how they dynamically exist within the human body. Within collagen, water is an integral part of its structure, where water migration from dehydration or osmotic stresses can lead to structural alterations such as shrinkage.^29^ It can be inferred that the state and amount of water in collagen, therefore, influences structural observations made at the nanoscale. In traditional TEM preparation, collagen fibrils or collagenous tissue sections are deposited onto TEM grids and then dried, either in air or alcohol-based schemes. This is one of the more affordable and accessible techniques for preparation, however, drying can lead to structural alterations arising from artifacts such as shrinking.^30^ In contrast, in cryo TEM preparation samples are flash-frozen and considered to be in a hydrated state. Thin frozen samples routinely lead to high-resolution (HR) imaging,^20,31^ but, preparing a cryo TEM sample has its technical challenges. Cryo preparation requires specialized equipment and samples can be susceptible to artifacts such as air-water interfaces and ice-induced crystals may influence the quality of samples produces and nanoscale observations,^32^ while the as-frozen state limits the ability to capture *in situ* dynamics.

Thus, capturing mineralization processes in a dynamic liquid state, without any alterations to the structure, is key to understanding and interpreting mineralization pathways. Liquid TEM, the room temperature correlate to cryo TEM, is a relatively new imaging technique that can provide HR and real-time information on hydrated biological states.^33–41^ While cryo TEM routinely provides HR imaging and is considered the gold standard for biological imaging, recent technical breakthroughs in acquisition strategies and thin film enclosures are pushing liquid TEM imaging resolution to comparable levels with the additional benefit of retaining the liquid state.^40^ Liquid imaging is often performed in one of two ways: with a commercial liquid TEM holder or by encapsulation methods. Both approaches have been used previously to probe simple CaP reactions, revealing intricate dynamic pathways behind the formation of apatite,^33,36,42,43^ however, neither approach has been used to observe collagen mineralization processes.

Herein, new liquid TEM techniques were used in a facile manner to probe collagen mineralization. A hybrid SiN-thin film enclosure was used with a simple two-step assembly process (Fig. 1), where collagen in solution was added to the enclosure, hermetically sealed, screened in a light microscope and loaded into the TEM. This method was highly repeatable – with an over 70% estimated encapsulation success rate, facile – taking only 10 mins to prepare, and affordable – accessible to use without specialized equipment.

**Figure 1.**
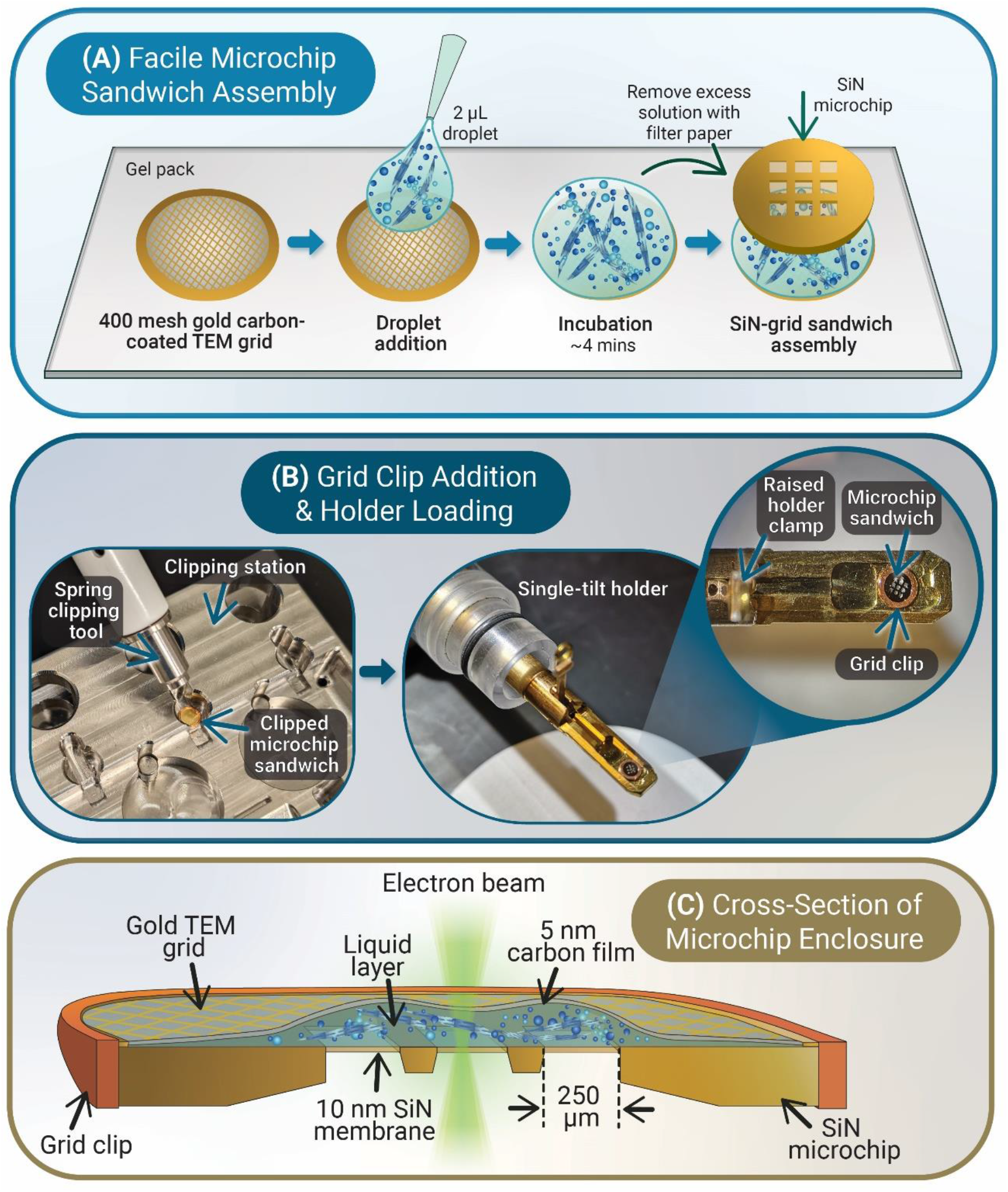
Microchip sandwich enclosure for liquid TEM of calcium phosphate-based collagen fibril mineralization processes. **A**. Assembly of liquid enclosure method from left to right, where 2μL of mineralization solution is deposited onto a plasma-cleaned carbon-coated 400 mesh gold TEM grid, incubated for 4 minutes, and then excess solution is removed before assembling the enclosure by adding a SiN microchip on top to create a SiN-grid sandwich. **B**. The SiN-grid sandwich enclosure is hermetically sealed using a cryo-autoloader grid clip, then stored or imaged immediately using a single-tilt TEM holder. **C**. Cross-section of the microchip enclosure, highlighting the thin film of the 5 nm carbon and 10 nm SiN membranes.

A rat tail tendon source was used to extract type I collagen, from which reconstituted collagen fibrils were prepared for mineralization and imaging, described in greater detail in the supplemental methods (Fig. S1). Type I collagen assembles into fibrils with a repeating periodicity of 67 nm, with characteristic gap and overlap zones of 40 nm and 27 nm, respectively.^8,11^ CaP-based solutions with pAsp were used to drive collagen mineralization, which has successfully mimicked intrafibrillar mineralization similar to bone *in vitro*.^16^ A 25 μg/ml concentration of pAsp (Mw 14 kDa; Alamanda Polymers, Huntsville, AL, USA) was used, where full mineralization of collagen fibrils (∼0.1 mg/ml) has been observed to occur within 18 hrs. Solutions were mineralized in bulk at 37°C and sampled using liquid TEM at time points of 4, 7, 15.5, and 18 hours, rather than mineralized within the enclosures in an attempt to mitigate long-term confinement effects described in literature.^44^ Automated low-dose image acquisition, automated focusing, frame averaging and direct electron detectors, detailed further in supplemental information, are some of the technological strategies implemented to enable imaging of the electron beam-sensitive samples.

Within a 4-hour mineralization period, two representative collagen stages were detected, with unmineralized regions dominating the mineralized (Fig. 2). Along less mineralized collagen fibrils both small and large higher-contrast globular particle aggregates were noted (Fig. 2A-B) and confirmed to be amorphous by SAED (Fig. 2B-Inset). This is indicative that the particles may be amorphous CaP (ACP) precursors, though elemental analysis would be necessary to confirm this. Mineral precursor aggregates have been identified in collagen models involving pAsp by works by Olszta et al.^18^ and Deshpande and Berniash^17^ with theories proposing a polymer-induced liquid precursor or involving particle attachment, respectively. Nudelman et al. theorize that collagen works together with pAsp to inhibit apatite formation and promote ACP infiltration observed in HR cryo TEM studies involving horse tendon-derived collagen.^20^ They note clusters of charged amino acids in gap and overlap zones form nucleation sites that facilitate ACP transformation into aligned mineral apatite platelets.^20^

**Figure 2.**
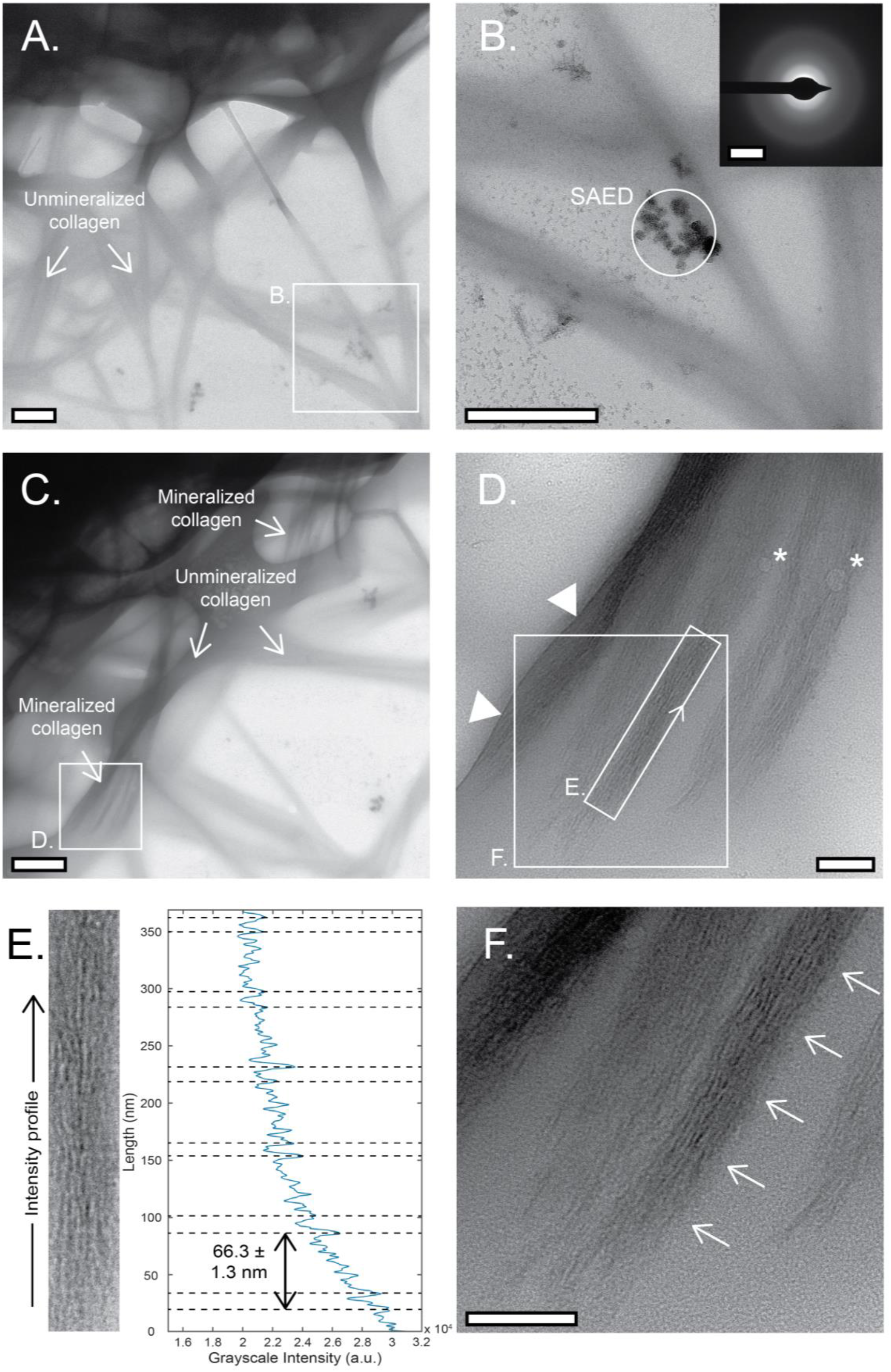
Representative BF TEM micrographs of hydrated mineralized collagen fibrils mineralized after 4 hrs in the presence of pAsp. **A**. Lower magnification image highlights highly mineralized (darker contrast) and nearby regions of unmineralized collagen fibrils (box) under hydrated conditions. **B**. Higher magnification image showing an intermediate phase of collagen mineralization where smaller mineral aggregates were seen in solution and attached to the fibrils. **Inset – B:** SAED of these mineral aggregates show that they are amorphous in nature. **C**. A separate low magnification region where sections of unmineralized and mineralized collagen fibrils are shown. **D**. Higher magnification region where mineral platelets were observed in the hydrated collagen fibrils, where a liquid waterfront (triangles) and bubbles (stars) validate the hydration of this region. **E**. Grayscale intensity plot profile from the rectangular box in D. shows faint collagen banding patterns that repeat at intervals of 66.3 ± 1.3 nm, highlighted visually in **F**. (arrows). Scale bars: A.-C. 500 nm, Inset-B. 5 nm^-1^, D. and F. 100 nm.

Within the 4-hour liquid TEM mineralization period probed, amorphous particles were observed present both adjacent and attached to collagen fibrils and did not appear to have liquid character. A caveat is that these observations cannot rule out the presence of crystalline-based and/or ion-dense liquid precursors elsewhere in the bulk sample. However, without the presence of collagen, the same CaP mineralization solutions combined with pAsp at 25 μg/ml resulted in the formation of amorphous precursor mineral phases in the form of branched polymeric assemblies or globular particle aggregates (Fig. S2). This observation aligns with assemblies visualized in similar CaP-pAsp-based systems, where works by Quan and Sone show that pAsp molecules increase crystallization inhibition effects by stabilizing ACP while recent *in situ* studies showed the formation of such assemblies through liquid TEM methods.^15,33,38^

In other regions at 4 hours, select fibrils show signs of more mature mineralization, with mineral platelets formed along fibril lengths (Fig. 2D-F). This is expected as the fibrils will not all mineralize simultaneously based on their heterogeneous distribution in suspension and variations in bundle sizes. The presence of mineral platelets at the 4-hour period demonstrates the active role of collagen in mineralization with pAsp, as the same solution without collagen at 4 hrs forms ACP polymeric branched structures as opposed to needle-like mineral platelets (Fig. S2). Faint collagen banding patterns that repeat locally at intervals of 66.3 ± 1.3 nm are shown in Fig. 2E, highlighted with better clarity in Fig. 2F (arrows). This local measurement is slightly lower than values obtained by Quan et al. reported as 68.0 ± 0.2 nm from cryo TEM of unmineralized collagen fibrils derived from rat tail tendon.^27^ This difference is expected due to contractile stresses that arise during mineralization and result in shrinkage.^27,29,45^

Similar trends were noted in correlative 4-hour collagen biomineralization study in dehydrated conditions (Fig. S3), where SAED results show the presence of a CaP apatite crystalline phase with diffraction pattern (DP) rings for planes (002), (211), and (004) observed (Fig. S3D-Inset). The (002) and (004) DP arcs in reciprocal space aligned with that of the collagen in real space, indicative of apatite *c-axis* alignment to the long axis of the fibril (Fig. S3D-Inset). However, even with sample rinsing, under dehydrated TEM preparation methods (Fig. S1C) salt residue was noted (Fig. S3A, B). Salt accumulation was not noted in hydrated conditions, however – pointing towards the importance of having a fully hydrated view of biomineralized samples.

At later time points, after the 7-hour mineralization period, similar regions with various stages of mineralization were observed (Fig. 3). In Fig. 3A, bubbles (upper right) exemplify hydration in densely packed collagen regions. Unlike many graphene liquid cells that feature isolated liquid pockets,^46^ continuous liquid layers in grid squares appear to be achieved. Figure 3B focuses on a collagen fibril that exhibits signs of early mineralization, surrounded by unmineralized fibrils seen at lower magnification (Fig. 3A). Particulate matter was also noted in solution and on collagen fibrils (Fig. 3B, triangles). To visualize the region with higher resolution, DED acquisition was also performed (Fig. 3C). Here, high-frame-rate 40 fps 4K videos were captured on the Direct-View camera at an electron dose rate of 45 e/Å^2^s (1-second exposure). Videos were frame averaged by processing with motion correction using cryoSPARC 2.14.2^47^ with MotionCor2 v1.2. DED acquisition shows, with high contrast, the presence of plate-like mineral features aligned along the collagen fibril among what appears to be amorphous darker contrast regions (Fig. 3C). SAED highlights faint diffraction patterns, the start of fine mineral platelets forming (Fig. 3C-Inset), though these are on a predominantly amorphous background without a distinguishable preferential orientation, as seen by images. Cryo TEM observations by Nudelman et al. suggest that collagen mineralization progresses by the infiltration of ACP and later the formation of apatite mineral platelets arises from ACP regions, with a similar intermediate crystalline platelet and amorphous regions noted.^20^

**Figure 3.**
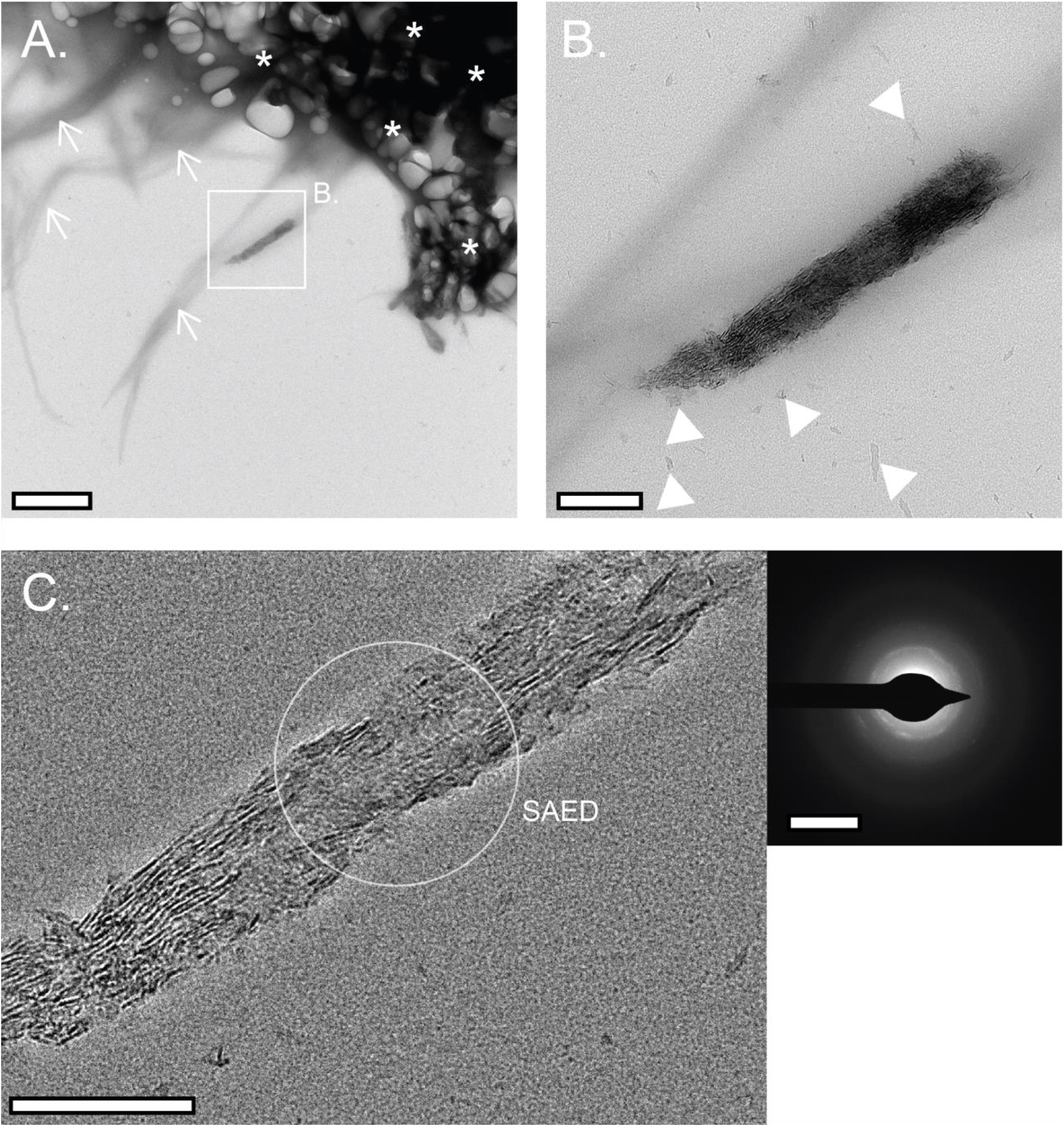
Representative BF TEM micrographs of hydrated mineralized collagen fibrils mineralized after 4 hours in the presence of pAsp. **A**. Low magnification imaging highlights a mineralized region (box) with nearby unmineralized fibrils (arrows) and thicker clusters of less well-dispersed fibrils (stars). Bubbles (in starred regions) indicate the imaging area is hydrated. **B**. Higher magnification image showing an intermediate phase of collagen mineralization wherein smaller mineral aggregates were seen in solution and attached to the featured fibril (arrowheads). **C**. DED acquisition (frame-averaged 4K 40 fps movie, 1-sec exposure) shows plate-like mineral features resolved clearly with high contrast in the hydrated fibril in select regions along its length. **Inset - C:** SAED shows the intermediate apatite crystallization phase wherein thin and faint arcs without preferred orientation are observed on an amorphous background. Scale bars: A. 1 µm, B. and C. 200 nm, Inset-C. 5 nm^-1^.

The fibril featured in Fig. 3B-C is only partially mineralized, with the transverse direction showing signs of mineralization across its width, while the longitudinal direction is mineralized only in the specific segment shown in Fig. 3B-C, though it extends further (Fig. 3A). This indicates active mineralization was ongoing before encapsulation for TEM imaging. The ends of this mineralized region seem partially mineralized, suggesting mineralization is still occurring at these mineralization fronts – progressing outwards, along the length of the fibril. This parallels the mineralization model proposed by Macías-Sánchez et al.^22^ whereby using a compact turkey-tendon collagen model, mineralization is said to occur first transversely across a fibril width and then proceeds longitudinally.

Nearly mature fully mineralized collagen is observed after longer times, with a representative period of 15.5 hours in Figure 4. Stable bubbles demonstrate the hydrated nature of these specimens (Fig. 4A). At a higher magnification region, discrete mineral platelets were observed aligned with their *c-axis* to the fibril length (Fig. 4B). Again, DED acquisition shows clear platelets aligned with collagen banding, suggesting the possibility of intrafibrillar mineralization (Fig. 4C). On the SAED, (002), (211), and (004) rings were noted with (002) and (004) arcs aligned to fibrils in real space (Fig. 4C-Inset1,2), confirming the presence of apatite with preferred *c-axis* along the fibril length. Imaging of the mature 18 hr collagen mineralization period (Fig. S4) further supports that the enclosure membranes and liquid environment have limited influence on interpreting SAED patterns.

**Figure 4.**
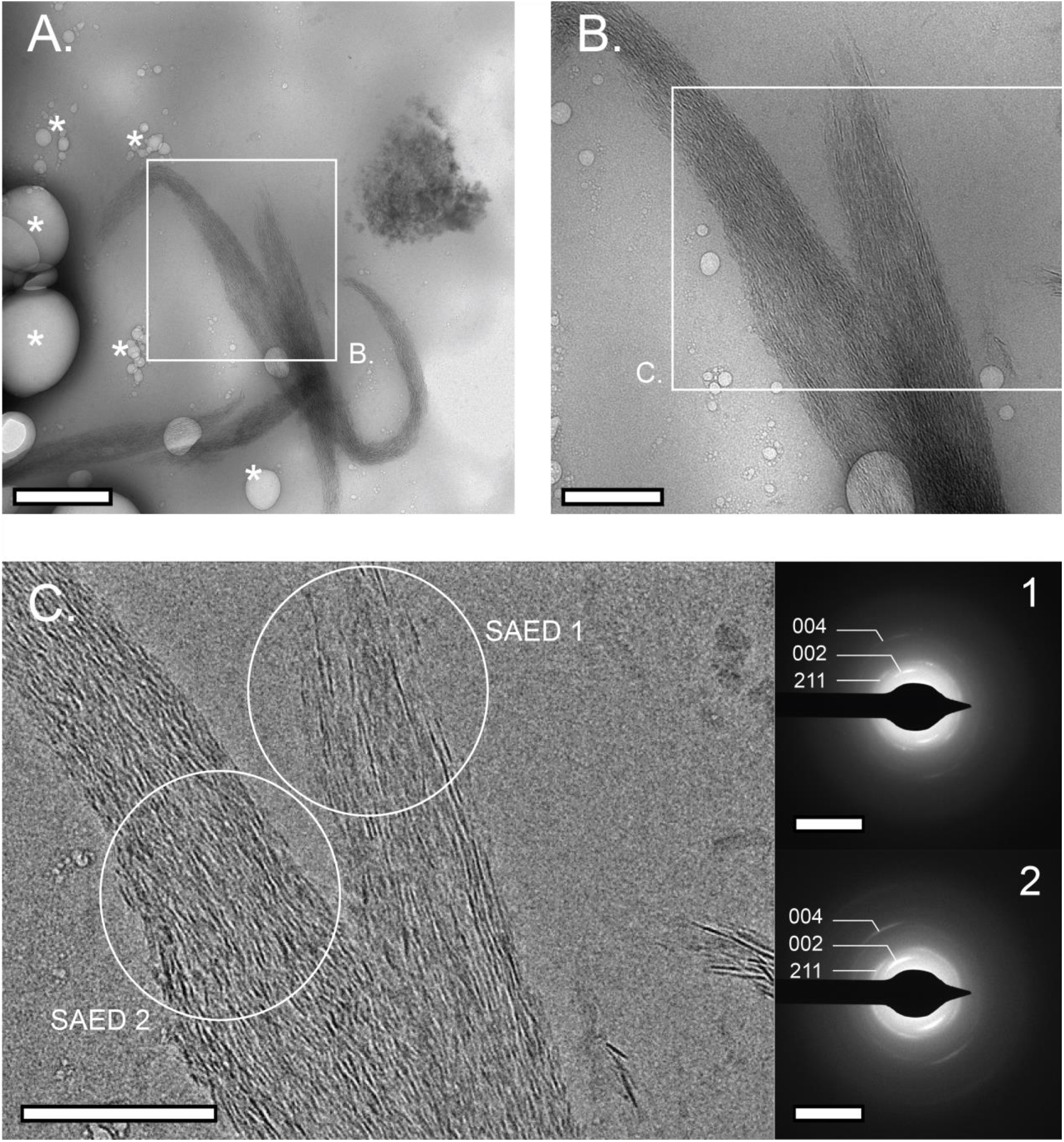
Representative BF TEM micrographs of hydrated mineralized collagen fibrils after 15.5 hours in the presence of pAsp. **A**. Low magnification imaging highlights visible mineralized collagen fibrils (box). Bubbles (stars) indicate the imaging area is hydrated. **B**. Higher magnification image shows a region with distinctive mineral platelets visualized to be aligned along the length of the collagen fibrils featured. **C**. DED acquisition (frame-averaged 4K 40 fps movie, 1-sec exposure) shows at higher resolution the clear definition of these platelets. **Insets 1**,**2 – C:** SAED shows 002), (211), and (004) rings corresponding to CaP-based apatite crystals. The (002) and (004) arc alignments are indicative that the crystalline apatite is preferentially oriented with its *c-axis* parallel to the long axis of the collagen fibrils. Scale bars: A. 500 nm, B. and C. 200 nm, Insets-C. 5 nm^-1^.

Collagen fibrils and their aggregates were challenging to image in liquid TEM. First, they are quite large for liquid TEM enclosures, with ranges in size from twenty to hundreds of nanometers in diameter, which also corresponds to sample thickness.^48^ Moreover, collagen fibrils distribute heterogeneously in solution and tend to aggregate. This poses a challenge for commercial liquid TEM holders that use rigid SiN membranes, where the large collagen fibrils would act as spacers – leading to thick liquid layers that limit the resolution and/or uneven membrane stresses that lead to rupture.^37^ While affordable graphene thin-film liquid TEM enclosures are now often used and result in thinner liquid layers for HR imaging,^49–54^ their assembly can be complex with multiple steps and the handling of fragile graphene susceptible to cracking.^51,55^ Moreover, only small amounts of liquid are captured within these assemblies, where liquid pockets enveloping samples are often achieved rather than continuous liquid layers.^55^ The effect of the lack of liquid on mineralization would likely be substantial; others have highlighted that liquid confinement can influence the stabilization of mineral precipitates^44^ and increase ACP size.^36^ The hybrid carbon-SiN membrane liquid TEM thin-film enclosure used herein (Fig. 1) proved to be effective in imaging these large samples with continuous liquid layers while automated tools and direct electron detector acquisition allowed for high-throughput HR imaging at low dose.

While this exciting new strategy holds promise for studying biomineralization reactions in hydrated conditions, a limitation was that dynamic events could not be captured with this imaging strategy. This is theorized due to the following three factors, discussed herein, which must be considered in the future: i) confinement effects, ii) lack of heating functionality of the enclosure, and iii) cumulative beam damage.

Firstly, based on qualitative changes observed in liquid contrast, it is likely that the carbon TEM-grid film ‘blankets’ the collagen structures, where near the fibrils the collagen itself acts as a local spacer but away from the fibrils the liquid thins out. Confinement of mineralized solutions in small liquid volumes is known to affect mineralization, namely by influencing nucleation rates and leading to larger ACP precursors in comparison to bulk volumes.^36,44^ By using a bulk mineralization solution and probing at defined time points, some long-term confinement effects may have been avoided in this work. Further, the thin liquid layers and encapsulation may have provided more of a 2D, rather than 3D, space for mineralization where collagen fibrils and large particles were immobilized within the enclosure. Similar observations have been made in carbon-based thin-film liquid cells of larger biological samples such as bacteria, which are described as being wrapped in the film with limited movement,^56^ while smaller samples, such as adeno-associated virus^40^ and platinum nanocrystal growth in solution,^57^ have been reported to exhibit dynamic motion in such enclosures.

Secondly, the current enclosure has no heating functionality to mimic physiological conditions in which collagen biomineralization occurs at 37°C. Thermodynamics play a key role in calcium phosphate mineral nucleation and particle growth rates, as summarized in detail by Sorokina et al.^58^ Energy transferred thermodynamically into a system by heat contribute to the control of nucleation and growth, affects the diffusion of mineralization products, and influences the thermodynamic stability of pre-nucleation and ACP phases present in the system.^58^ Considering the colder vacuum TEM environment, this may have delayed mineralization within the enclosures, nearly halting reactions from progressing and limiting dynamics.

Last of all, the electron beam sensitivity of the solution and cumulative beam dose damage contribute to challenges related to capturing dynamic mineralization processes. In this mineralization system, the use of buffers stabilized to a pH of 7.4 is hypothesized to aid in reducing chemically active species^33^ while in mineralized fibrils apatite acts as a scavenger of radiolysis species to increase electron beam dose damage thresholds.^59^ However, in thicker liquid regions where small particle motion was observed during screening, beam sensitivity of the solution caused near-instant bubbling (Fig. S5A). Such bubbles form from H+ ions arising from beam irradiation during imaging of beam-sensitive solutions and in thick regions that require a higher beam intensity to resolve formed products.^59,60^ Automated imaging through SerialEM helped overcome these barriers to mitigate beam exposure during sample screening, though beam-sensitive regions still posed problems for longer video acquisitions needed to capture dynamics. Cumulative doses of approximately 250 e/Å^2^ were detected to lead to severe structural beam damage in mineralized fibrils (Fig. S5B). Moser et al.^60^ report that structural damage such as shrinking occurs in biological cell samples as low as 4 e/Å^2^ while structural damage caused by bubbling in cryo TEM occurred in ranges between 300-400 e/Å^2^.

## CONCLUSION

This work demonstrates a facile method to visualize collagen mineralization in liquid TEM for the first time. A new hybrid SiN-thin film liquid TEM enclosure hermetically sealed with a clip was presented which encapsulated collagen fibrils in thin, continuous liquid layers for imaging. Automated acquisition and direct electron detection were attributed to high-throughput, low-dose, HR imaging of collagen in liquid. In addition, automated imaging helped in increasing acquisition efficiency in screening samples and identifying regions of interest, while mitigating unnecessary electron beam exposure prior to acquisition.

Using these imaging techniques, various stages of mineralization were captured in liquid: an amorphous mineral phase was noted attached to unmineralized fibrils at initial time points, then amorphous and crystalline mineral platelets co-existed, and finally, distinctive nanoscale mineral platelets that aligned with their *c-axis* along the collagen fibrils were revealed and confirmed through SAED to be a crystalline apatite. These observations parallel and complement the findings of others under dry or cryo conditions,^9,11,20,22–27^ validating this liquid TEM technique as a feasible tool to apply to the investigation of complex real-time mineralization processes. This work highlights a significant leap forward by introducing a facile liquid TEM characterization tool to view native hydrated specimens with nanoscale features – important for understanding collagen mineralization and foundational to the clinical study of mineralized tissues and their repair.

## ASSOCIATED CONTENT

## SUPPORTING INFORMATION

The following supporting information is available free of charge:

- Supporting Information document containing Methods and Methods section as well as Supplemental Figures (PDF format).

## Notes

The authors declare no competing financial interests and no conflicts of interest.

## Author Contribution

**Conceptualization:** Liza-Anastasia DiCecco, Ruixin Gao, Eli Sone, Kathryn Grandfield

**Methodology:** Liza-Anastasia DiCecco, Ruixin Gao, Deborah F. Kelly

**Data curation:** Liza-Anastasia DiCecco, Deborah F. Kelly

**Formal analysis:** Liza-Anastasia DiCecco, Ruixin Gao, Deborah F. Kelly, Eli Sone, Kathryn Grandfield

**Funding acquisitions and supervision:** Deborah F. Kelly, Eli Sone, Kathryn Grandfield

**Writing:** Liza-Anastasia DiCecco, Eli Sone, Kathryn Grandfield

**Review and editing:** Liza-Anastasia DiCecco, Ruixin Gao, Deborah F. Kelly, Eli Sone, Kathryn Grandfield

## ACKNOWLEDGEMENTS

Financial support is greatly acknowledged from the Canadian Institutes of Health Research Project Grant (to E.D.S. and K.G. grant no. PJT 180575), and several programs of the Natural Sciences and Engineering Research Council of Canada (NSERC): Discovery Grant Program (to K.G. grant no. RGPIN-2020-05722), Canada Research Chairs Program (to K.G.; Tier II Chair in Microscopy of Biomaterials and Biointerfaces), and Vanier Canada Graduate Scholarship (to L.-A.D.). Electron microscopy was performed at the Canadian Centre for Electron Microscopy and the Pennsylvania State University Materials Research Centre and Huck Institutes of the Life Sciences. Carol Barter from Pennsylvania State University is kindly acknowledged for their electron microscopy technical support and guidance.

