## Supplemental Materials for "Liquid Transmission Electron Microscopy for Probing Collagen Biomineralization"

### 1. MATERIALS AND METHODS

#### i. Collagen Fibrillogenesis

Collagen was extracted from a rat-tail tendon model as a close structural mimic to type I collagen found in mineralized tissues within the human body.[1,2] Collagen fibrillogenesis and crosslinking of this model are further elaborated in works by Lausch et al.[3] Briefly, extracted rat tail tendons were dissolved at 1 mg/mL in 0.5 M acetic acid. Then, the solution was diluted 1:0.5 by distilled water (di$H_2O$) and the pH was adjusted to 5.0 with 1 M NaOH, which was incubated for 2 hours in a water bath at 35°C before collecting collagen fibrils.

Collected collagen fibrils were fixed using glutaraldehyde over an 18-hr fixation period with 0.08% glutaraldehyde in an aqueous solution (Electron Microscopy Sciences, Hatfield, PA, USA). Samples were subsequently dialyzed against di$H_2O$ for 2 days with frequent water replacements to halt system reactions and remove glutaraldehyde and acetic acid from the system.

#### ii. Collagen Mineralization

An overview of key experimental details within this and the following subsections of the materials and methods is provided in Fig. S1. Reconstituted collagen fibrils were mineralized based on procedures adapted from published work by the team of Despande and Beniash[4], where a calcium phosphate-based system is used. Mineralization was prepared by mixing equal volumes of collagen fibrils in $diH_2O$ at a concentration of approximately 0.4 mg/mL with the following aqueous-based solutions, all buffered to a pH of 7.4 at 37˚C using 1 M of hydrochloric acid: 9 mM $CaCl_2$ in 50 mM Tris, 4.2 mM $K_2HPO_4$ in 50 mM Tris, and 100 mM Tris–500 mM NaCl (Tris-NaCl). Poly-L-aspartic acid (pAsp) sodium salt (pAsp 100 – Mw 14 kDa; Alamanda Polymers, Huntsville, AL, USA) was dissolved into the Tris-NaCl solution such that a desired concentration of 25 µg/ml of pAsp would be achieved in the final solution. All mineralization solutions, except the collagen suspension, were filtered through a 0.2 µm acrodisc syringe filter before incorporation. After mixing, the final mineralization solution consisted of 125 mM NaCl, 1.7 mM $CaCl_2$, 9 mM $Na_2HPO_4$, 50 mM Tris, and a concentration of 25 µg/ml of pAsp and an approximate concentration of 0.1 mg/mL of collagen fibrils. Prepared solutions were mineralized at 37˚C in a water bath set with a probe-controlled hot plate and gentle mixing of 100 rpm. All reagents described were purchased from Sigma Aldrich and dissolved in Milli-Q $diH_2O$, except for the pAsp which was purchased Alamanda Polymers as noted.

For liquid TEM, mineralization periods of 4, 7, 15.5, and 18 hrs were probed to capture early to mature mineralization phases within collagen fibrils. For correlative studies of the mineralization solution without collagen fibrils featured in the supplemental material, the solution was prepared in the same manner with simply $diH_2O$ added instead of the collagen fibril suspension.

#### iii. Liquid Enclosure TEM: Sample Preparation and Acquisition

Figure 1 of the main body of this article highlights schematically the two-step assembly process of the liquid enclosure used in this work, adapted from the protocol described in DiCecco et al.[5] The hybrid SiN-thin film enclosure used is composed of two pieces: a 2.9 mm diameter silicon nitride microchip with nine membrane windows (sizes: (8) 250x250 µm + (1) 250x500 µm) with a membrane thickness of 10 nm (SiMPore Inc., West Henrietta, NY, USA) on one side and a gold 400 mesh TEM grid with a 5 nm carbon film (Electron Microscopy Sciences) on the other, clamped together by copper grid clip used for cryo-TEM autoloaders (ThermoFisher Scientific, MA, USA). Before loading, all TEM grids and SiN microchips were plasma glow discharged for 2 cycles of 45 seconds with a plasma current of 15 mA using a Pelco EasiGlow instrument (Ted Pella, Inc. Redding, CA, USA).

To form the enclosure, 2 µL of the liquid collagen solution was deposited onto the carbon side of a TEM grid resting on a gel pack holder (Fig. 1A). The grid was then incubated for 4 minutes to allow collagen to settle. During this time, the samples were screened under a light microscope to confirm collagen deposition. Following this, excess solution was removed with the edge of a piece of Whatman #1 filter paper (Sigma-Aldrich Canada Co., Oakville, ON, Canada), and the sample was enclosed by the addition of the SiN microchip (Fig. 1A) on top. Early attempts at collagen deposition onto SiN highlighted that collagen removed itself easily from the membranes, while the carbon-coated TEM grid kept samples stable on the surface and was thus used. The assembly was hermetically sealed with a copper grid clip, traditionally used for cryo TEM autoloaders (Fig. 1B). Prior to insertion in the TEM, the enclosure was inspected using a light microscope to ensure there were no cracks in the SiN microchip and that collagen fibrils were visible.

For correlative dry TEM samples presented within the supplemental of this study, 2 µL of a liquid mineralization sample was deposited onto a copper 400 mesh TEM grid with either a continuous or lacy 5 nm carbon film (Electron Microscopy Sciences) for reactions with or without collagen, respectively. These samples were then incubated for 4 minutes and then dipped into methanol to halt any further mineralization reactions and rinse away excess products arising from drying.

A single-tilt TEM holder was used for bright field (BF) TEM imaging using a Talos F200C or X series TEM (ThermoFisher Scientific, MA, USA), both operated at a voltage of 200 kV and equipped with a Direct-View direct electron detector (Direct Electron, CA, USA) and/or a CETA camera. For liquid TEM, the SerialEM software[6] (University of Colorado, Boulder, CO, USA) was used to facilitate high-throughput acquisition through automation using both the CETA camera and Direct-View detector. [5]Automated sample screening using SerialEM provided low-dose, low-magnification window-membrane maps, which supported quick sample screening and the selections of regions of interest (ROIs) for imaging. This offered quick screening capabilities of the entire microchip sandwich for ROIs, limiting beam exposure to the sensitive samples. Within each TEM grid square captured, ROIs for imaging were either selected manually or chosen through a grid square of points for serial acquisition, where SerialEM automated focusing was used to adjust acquisition parameters adjacent to an ROI to mitigate its beam exposure to an ROI.

Electron dosage was varied between 40-55 e/$Å^2$ per acquisition. When considered, direct electron (DE) movies were acquired at 40 fps, with 1 s exposure at an approximate rate of 45 e/$Å^2$s. These DE movies were frame averaged to provide images featured within this work with motion correction using cryoSPARC 2.14.2[7] with MotionCor2 v1.2.3. Selective area electron diffraction (SAED) was achieved using either 10 µm or 40 µm sized apertures which covered an approximate diameter of 125 nm or 250 nm in the real image, respectively. Indexing for SAED polycrystalline patterns for CaP-apatite was done by analyzing concentric ring patterns belonging to the polycrystalline structures captured using the ImageJ software. From these measurements of the calibrated camera acquisitions, interplanar spacing was calculated and compared to reference CaP-based crystal structures. Regions with distinguishable ring patterns were compared to a hydroxyapatite crystallography base reference[8] – as expected for mineral platelets present in mineralized collagen fibrils within regions investigated.

**iv. Line Scan Analysis**

When referenced, the periodicity of collagen fibrils was investigated using grayscale intensity analysis using the ImageJ software package “Plot Profile” function. Here, the fibril considered is featured in Figure 2 with 6 distinguishable periods noted. This region was considered for line grayscale intensity profiling along the c-axis of the collagen fibril, with an applied averaging line width of approximately 47.7 nm (300 px). Local graph maximum and minimum points were identified using MATLAB, where a custom smoothing algorithm[9] was applied for local max/min peak identification using pseudo-Gaussian smoothing (4 passes of the same sliding average) applied to 5 data points at a time. The periodicity of the collagen fibril was calculated based on the appearance of a local maximum present, visually associated with collagen banding patterns, found at repeating patterns of the grayscale intensity plots.

### 2. SUPPLEMENTAL IMAGES

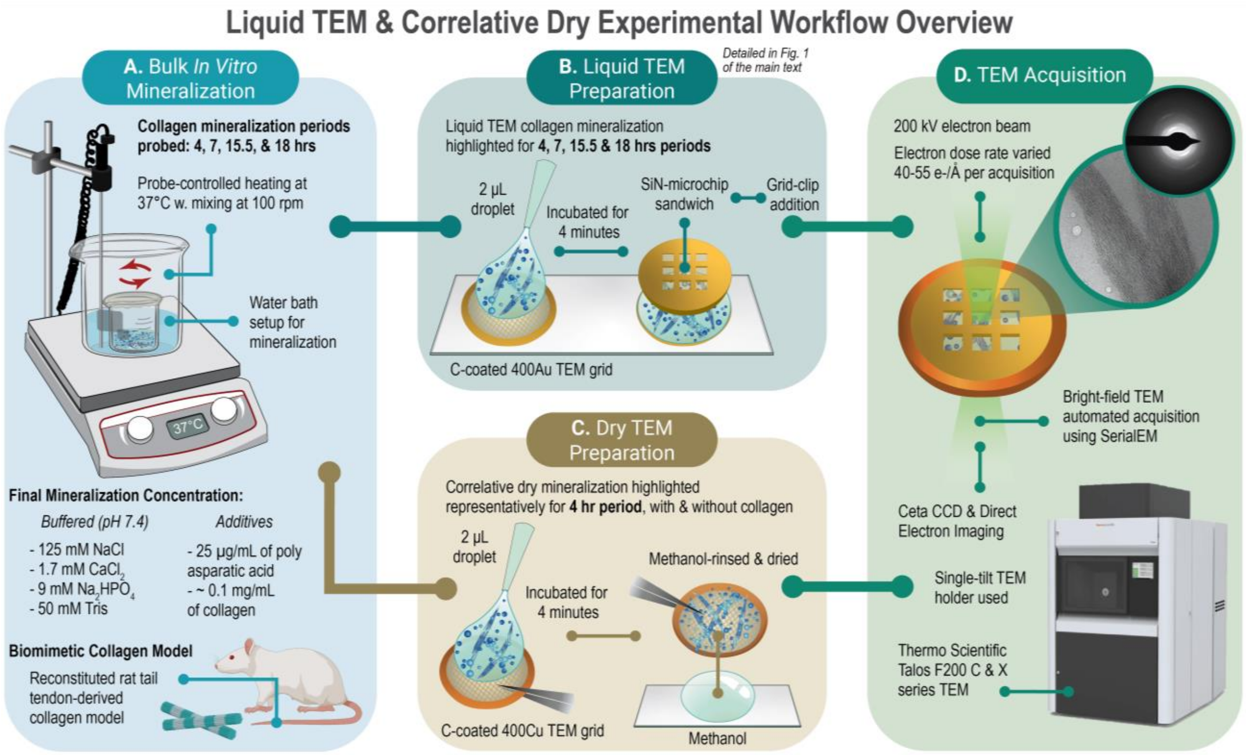


**Figure S1. Liquid TEM and Correlative Dry Experimental Workflow Overview. A.** Summary of mineralization conditions and setup considered for the study, where collagen mineralization periods of 4, 7, 15.5, and 18 hrs were considered for liquid TEM. **B.** Summary of liquid TEM sandwich enclosure preparation, explained in greater detail within Fig. 1 of the main text. **C.** Correlative dry TEM preparation method highlighted for the supplemental results featured with and without collagen for the 4-hr mineralization period. **D.** Overview of key TEM acquisition parameters. The murine schematic in A. was created with Biorender.com.

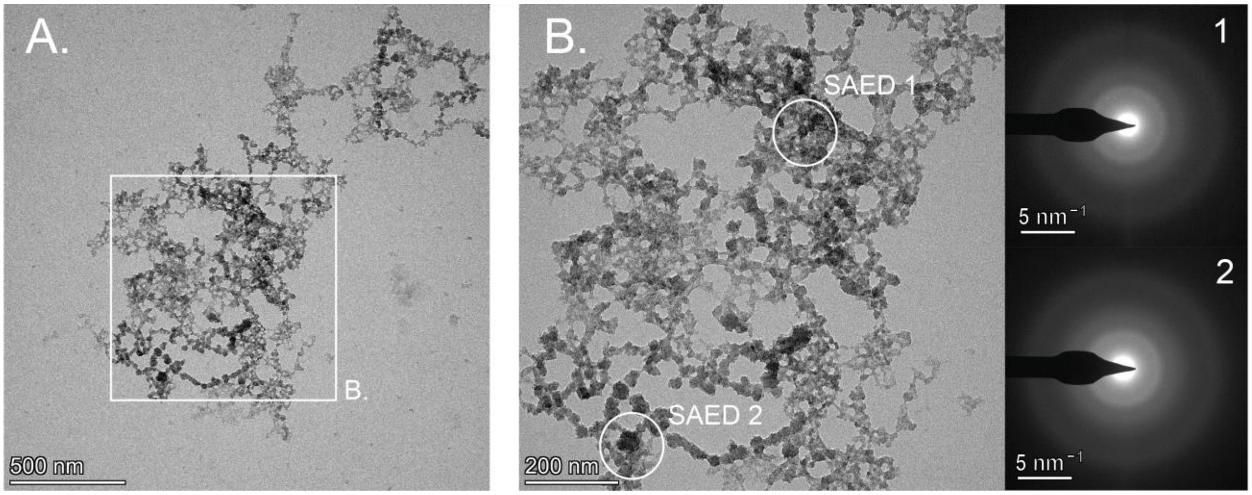


**Figure S2. Representative correlative *ex situ* BF TEM micrographs of dry CaP products mineralized after 4 hrs in the presence of pAsp without collagen. A.** Lower magnification image highlights larger mineral aggregates that formed within the solution. **B.** Higher magnification image shows branched spherical mineral assemblies. **Insets 1, 2 – B:** SAED of regions marked in B. show the amorphous nature of mineral aggregates.

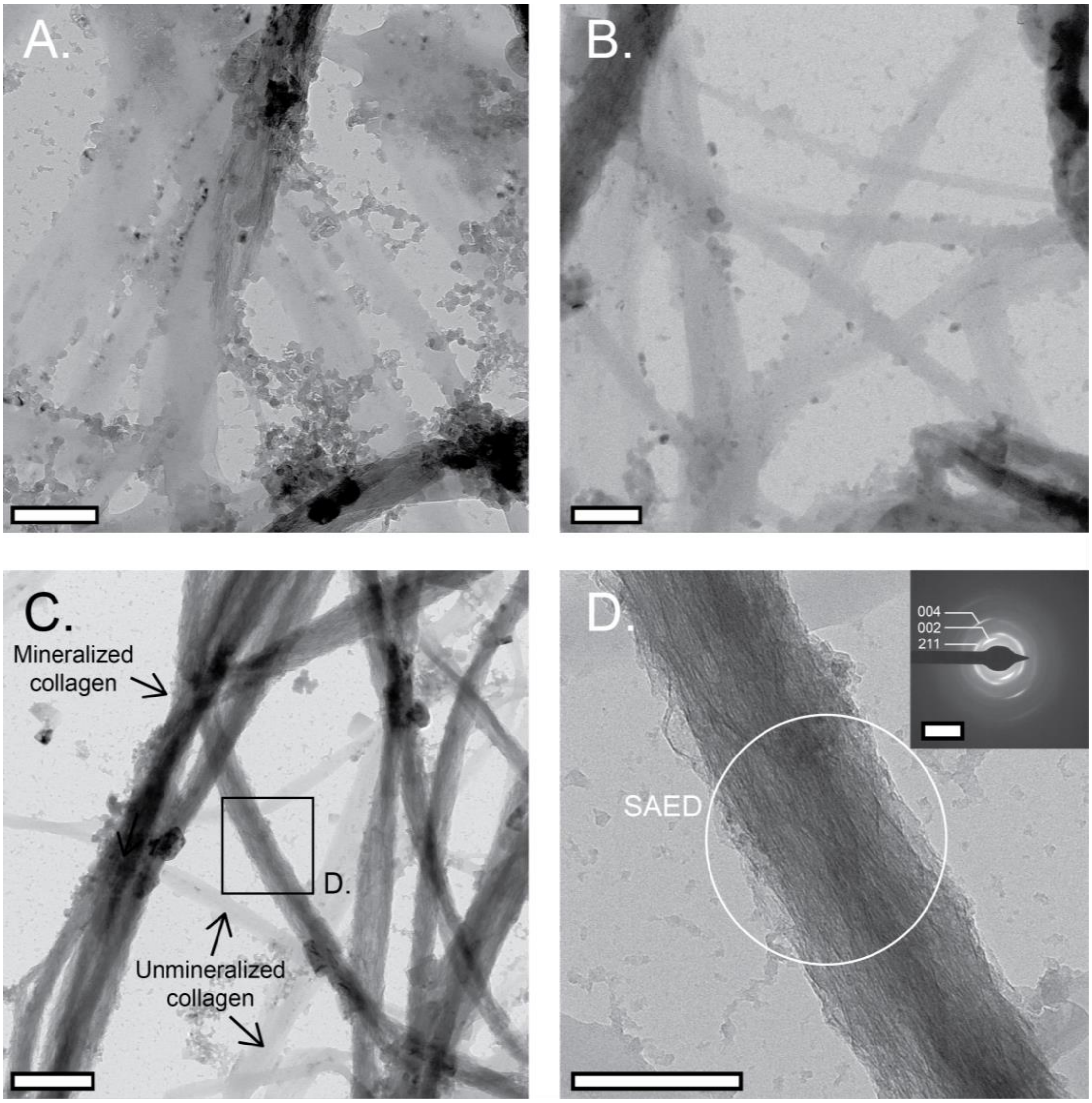


**Figure S3. Representative correlative BF TEM micrographs of dry mineralized collagen fibrils after 4 hrs in the presence of pAsp. A.-B.** Representative regions consisting primarily of unmineralized fibrils at an intermediate phase of collagen mineralization where small mineral aggregates can be seen around and attached to the fibrils. Despite rinsing, salt residue from mineralization buffers is noted to possibly adhere to the fibrils – a drawback to this traditional dry-based preparation method. **C.** Another region highlights an area in which both unmineralized and mineralized collagen fibrils can be seen **D.** Higher magnification region where mineral can be observed along the length of the collagen fibril. **Inset – D:** SAED show the (002), (211), and (004) rings corresponding to CaP-based apatite crystals, where (002) and (004) arcs alignment indicate that these mineral platelets are preferentially oriented with its *c-axis* parallel to the long axis of the collagen fibrils. Scale bars: A., B., and D. 200 nm, C. 500 nm, Inset-D. 5 $nm^{-1}$.

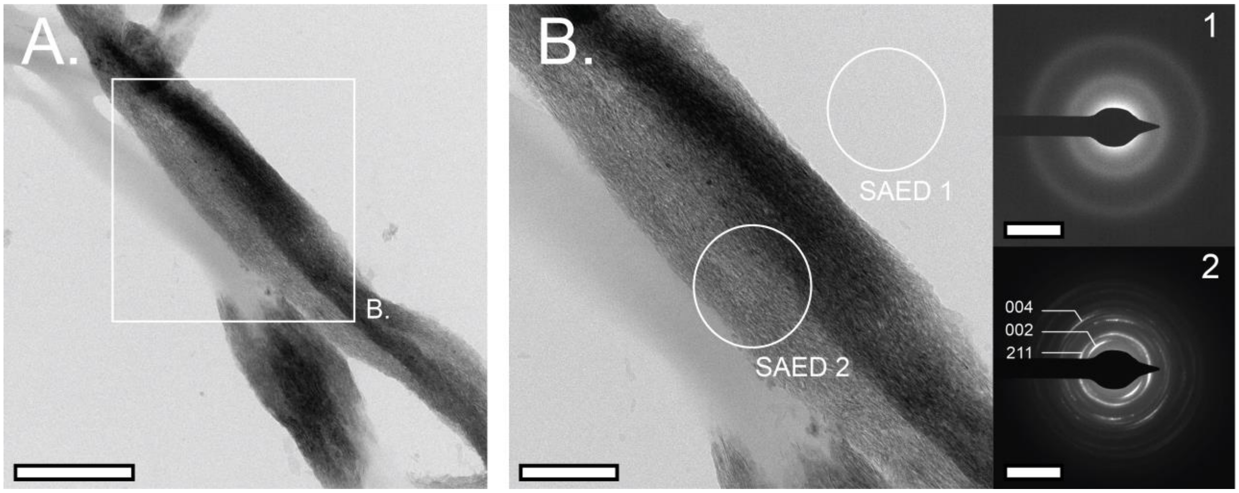


**Figure S4. Representative BF TEM micrographs of hydrated mineralized collagen fibrils mineralized for 18 hrs, showing minimal influence of the enclosure on SAED interpretation. A.-B.** Low to high magnification imaging highlights highly mineralized collagen fibrils, with mineral (darker contrast) appearing along the fibril length. **Inset 1– B:** SAED of the substrate in the region without collagen shows the amorphous nature of the SiN and carbon film membranes. **Inset 2 – B:** SAED show the (002), (211), and (004) rings corresponding to CaP-based apatite crystals, where (002) and (004) arcs alignment indicate that these mineral platelets are preferentially oriented with its *c-axis* parallel to the long axis of the collagen fibrils. These patterns are noted clearly within the region, despite the apparent thickness of this collagen fibril bundle, indicating that the SiN and carbon film membranes are thin enough to obtain clear diffraction patterns of hydrated specimens with limited influence on signal detection. Scale bars: A. 500 nm, B. 200 nm, Insets-B. 5 $nm^{-1}$.

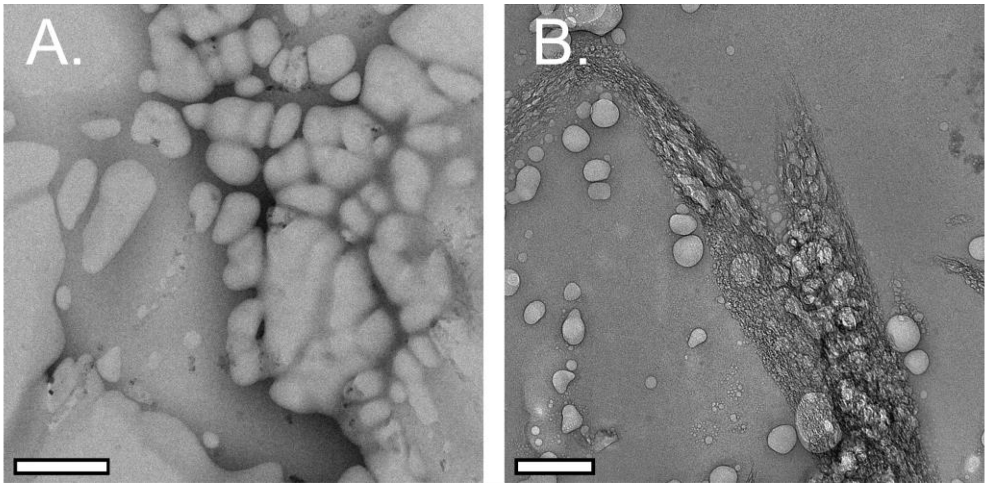


**Figure S5. Representative regions of thicker liquid and bubbling artifacts in 15.5 hr collagen mineralization sample. A.** BF TEM micrograph of thicker liquid region showing instantaneous bubbling on imaging of the solution, highlighting challenges with beam sensitivity and related resolution limitation caused by liquid mobility within thick regions in this liquid TEM imaging. **B.** Overview image of the region of interest featured in Figure 4 of the main text after continuous imaging and severe beam damage, occurring after a cumulative electron dose of approximately 250 e/Å$^2$. Scale bars: A. 500 nm. B. 200 nm.
